# Compact Vision–Language Models Enable Efficient and Interpretable Automated OCT Analysis Through Layer Specific Multimodal Learning

**DOI:** 10.1101/2025.08.07.669187

**Authors:** Tania Haghighi, Sina Gholami, Jared Todd Sokol, Jennifer I. Lim, Theodore Leng, Atalie C. Thompson, Hamed Tabkhi, Minhaj Nur Alam

## Abstract

Translating the intricate anatomical signatures of retinal disease from OCT B-scans into clear, accurate clinical narratives demands AI models that seamlessly fuse visual features with domain expertise. We curated a multimodal dataset of 40,000 OCT B-scans from public repositories and private clinical cohorts, each paired with expert validated summaries spanning six conditions: diabetic macular edema, diabetic retinopathy, geographic atrophy, drusen, choroidal neovascularization, and healthy retina. We introduce LO-VLM, a compact (247M parameter) vision–language model (VLM) that infuses anatomical guidance into both encoder and decoder for free form summary generation and multiclass disease classification. Benchmarking against state-of-the-art RetinaVLM, LLaVA-Med, and a ViT vision only model demonstrates superior performance. In a blinded evaluation by three board certified retina specialists scored the generated summaries, LO-VLM narratives achieved mean = 8.5 (standard deviation = 1.15) out of 10, compared to mean = 5.5 (standard deviation = 1.13) for RetinaVLM (p < 0.0001). In quantitative evaluations, LO-VLM achieved an SBERT similarity of 0.803 and a BERTScore F1 of 0.715, representing improvements of 8.2% and 28.8% over specialized VLM baselines. For disease classification, LO-VLM reached 96% accuracy (F1 = 96%), outperforming ViT by 13% and exceeding medical VLM benchmarks by over 62%. By reconciling interpretability with computational efficiency, LO-VLM establishes a new paradigm for efficient AI models in OCT interpretation.

## Introduction

Medical imaging and artificial intelligence (AI) tools have become indispensable in ophthalmic diagnostics. Recent successes in machine learning (ML) and deep learning (DL), coupled with the advent of unimodal and multimodal foundational models, have demonstrated expert level performance in tasks ranging from lesion detection to anatomical segmentation [1–4]. In the current clinical setting, optical coherence tomography (OCT) [5] has become the definitive modality for non-invasive retinal imaging, providing cross-sectional views of retinal microstructures with micrometer-scale resolution [6]. By employing low-coherence interferometry, OCT enables visualization of individual layers in vivo. These high resolution images are indispensable for diagnosing and monitoring ocular pathologies, including age-related macular degeneration (AMD), diabetic retinopathy (DR), and glaucoma [7–9]. Despite high fidelity for retinal disease diagnosis, manual assessment of OCT volumes can be both labor intensive and susceptible to variability between observers. Two clinicians may produce divergent measurements of retinal thickness or disagree on the presence of subtle hyperreflective changes. Moreover, the unequal distribution of trained retina specialists in rural or resource limited regions exacerbates diagnostic delays and potentially compromises patient outcomes. Automated tools that can pre-screen OCT scans, quantify relevant biomarkers, or prioritize cases for specialist review are therefore critical to optimizing clinical workflows and expanding access to timely eye care [10–12].

To meet this demand, recent advances in DL have demonstrated considerable promise for OCT analysis. Convolutional neural networks (CNNs) have been employed to classify OCT volumes according to disease state, distinguishing for example between normal retinas and those exhibiting signs of diabetic macular edema (DME) [13, 14]. CNNs have also been used to segment retinal layers in order to derive quantitative metrics such as central retinal thickness. These methods rely exclusively on image based feature extraction [15–18]. Despite achieving high performance on benchmark datasets, most existing approaches produce only a categorical diagnosis or a delineation of anatomical boundaries.

Notably, The ability to generate textual descriptions of retinal abnormalities similar to a narrative dictated by an expert clinician offers several advantages. Natural language summaries render AI outputs more interpretable and actionable [19–21]. Text based outputs can also be integrated directly into electronic medical record systems, streamlining documentation and reducing transcription errors. In regions with limited specialist availability, a model capable of producing reliable, human readable interpretations may serve as a decision support tool for general practitioners or optometrists. Therefore, there has been a growing interest in vision-language models (VLMs) pre-trained on extensive image–text corpora. By employing cross-modal pre-training strategies, these frameworks jointly optimize visual and linguistic encoders, aligning image features and textual semantics within a unified space. Such integrated representations can enhance diagnostic performance and facilitate the generation of concise, domain specific reports that are readily interpretable by clinicians. In recent years, researchers have demonstrated the utility of VLMs in several medical domains, especially in radiological applications [22, 23]. Building upon cross-modal pre-training strategies, ophthalmic VLMs such as RetinaVLM and VisionUnite [3] have been developed to generate narrative interpretations of retinal images. However, the considerable computational burden and limited pathological scope of these models underscore the necessity for an OCT-specific ophthalmic VLM that combines computational efficiency with the capacity to characterize the full spectrum of retinal pathologies in OCT imaging.

In this project, we present Layer-wise OCT VLM (LO-VLM), a domain specific VLM that employs layer specific multimodal learning to achieve efficient and interpretable OCT analysis for retinal disease diagnosis. LO-VLM jointly trains an image encoder and a text decoder optimized for accurately classifying retinal pathologies and quantifying essential biomarkers at individual retinal layers, all within a lightweight, computationally efficient framework tailored to the unique characteristics of OCT imaging. We address the dual challenges of ophthalmic multimodal data scarcity and computational efficiency in automated OCT interpretation through three key contributions:

- Creation of a Comprehensive Layer Specific OCT Image-Text Dataset: we construct a comprehensive dataset of 40,000 OCT image-text pairs featuring layer specific anatomical summaries that enable detailed retinal structure analysis and pathological characterization. This dataset addresses the critical gap in paired multi-modal data for OCT interpretation and provides a foundation for training VLMs in this specialized domain.
- Development of a Compact VLM for Multi-class Retinal Disease Diagnosis: we show that our compact VLM is capable of generating multi-class diagnoses by integrating both image and text inputs, rather than relying solely on image based analysis during training. We achieve high diagnostic accuracy across several disease categories, compared to image only model.
- Generation of Clinically Interpretable Layer Specific Descriptions: we demonstrate that our adapted model produces clinically meaningful layer specific anatomical descriptions that enhance interpretability and facilitate clinician validation of automated diagnosis. Through extensive quantitative and qualitative evaluations via clinician feedback, we show that the generated summaries align closely with expert annotations and support transparent human-readable reporting.

Central to LO-VLM is a layer-wise prompting strategy that injects explicit anatomical priors into the vision–language alignment stage. By forcing the model to attend to specific retinal layers, we (i) reduce semantic ambiguity and label noise, (ii) encourage fine-grained cross modal alignment at clinically meaningful loci, and (iii) achieve comparable representational power with a 247M parameter BLIP backbone instead of multi-billion parameter generic/medical VLMs. This design choice explains LO-VLM’s superior data efficiency and interpretability beyond raw accuracy gains.

## Results

This study establishes the effectiveness of layer specific multimodal training in LO-VLM, a compact 247M parameter VLM for OCT interpretation (Figure 1). By jointly optimizing a vision encoder and a text decoder on a curated dataset of 40,000 expert-reviewed, layer-specific image–text pairs and benchmarking against a ViT-Base baseline and three leading VLMs (RetinaVLM, LLaVA-Med, PaLI-Gamma 2), we show that LO-VLM: (i) achieves 96% disease classification accuracy — an absolute gain of 13% over ViT-Base (ii) attains state-of-the-art semantic similarity (SBERT = 0.803; BERT-F1 = 0.715) (iii) secures superior clinician ratings (mean 8.49 ± 0.88 vs. 5.41 ± 0.89; P < 0.0001) (iv) produces anatomically aligned saliency maps, and (v) maintains > 80% accuracy with as few as 248 training examples, demonstrating both robust interpretability and remarkable data efficiency.

**Figure 1:**
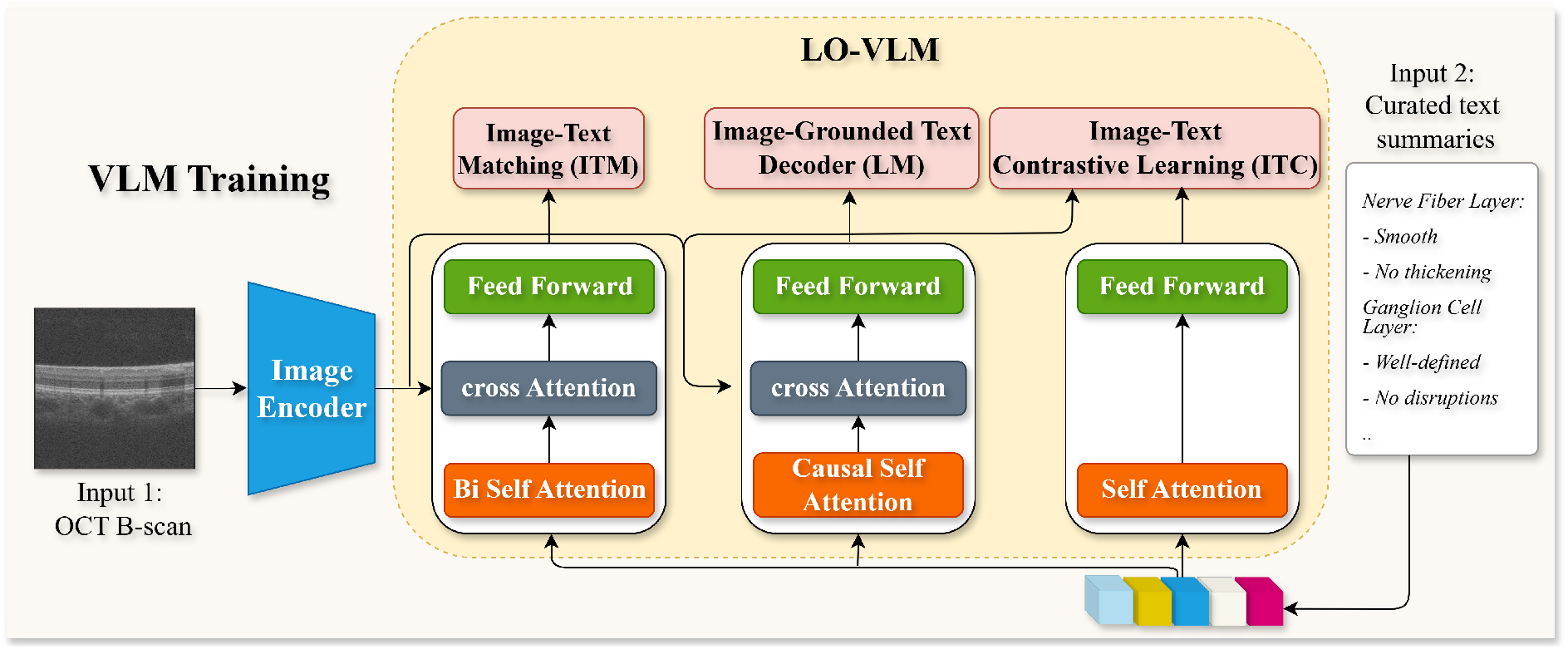
LO-VLM framework: visual embeddings extracted from OCT B-scans are jointly optimized with curated, layer specific text summaries via three complementary objectives.

### Baseline Comparisons

We began our evaluation by benchmarking three state-of-the-art VLMs on our held-out 1,000 test set. Specifically, we evaluated: (i) RetinaVLM, the only OCT specialized summary generator; (ii) LLaVAMed, a broadly pre-trained medical VLM with demonstrated OCT summarization skills (Figure 2; and (iii) PaLI-Gemma 2, a 3 billion parameter general purpose VLM trained on our 39,000 image–caption pairs. This structured comparison allowed us to untangle the contributions of domain specificity, medical pre-training, and sheer model scale to automated OCT interpretation.

**Figure 2:**
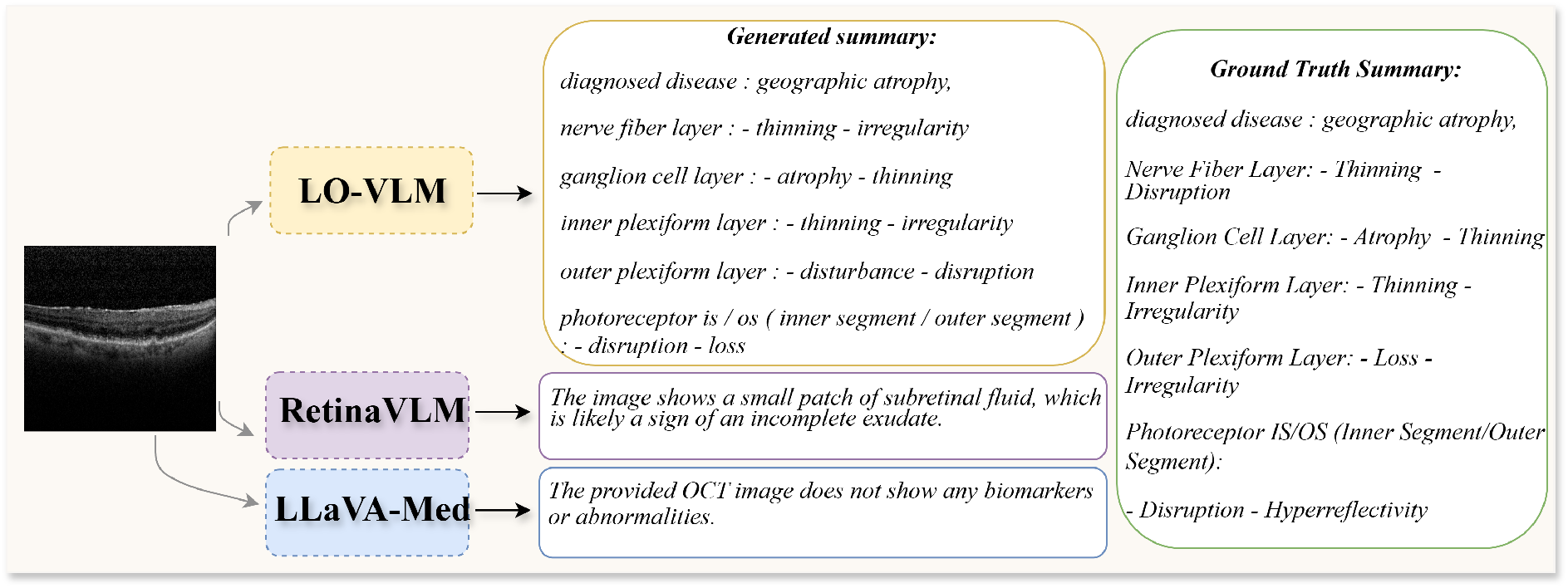
compares the outputs of LO-VLM, RetinaVLM and LLaVA-Med on the same OCT B-scan. For these comparisons, LLaVA-Med was prompted with “###Human: What biomarkers or abnormalities, if any, are present in the provided OCT image? ###Assistant:”, and RetinaVLM was prompted with “Describe the OCT image focusing on Biomarkers and abnormalities.”

To understand just how much textual grounding adds, we then trained a purely visual ViT-Base classifier on those same 39,000 OCT B-scans and evaluated it on the six classes diagnostic task: Normal, Drusen, Choroidal Neovascularization (CNV), DME, DR, Geographic Atrophy (GA). By directly contrasting its performance against the three VLMs, we could quantify the concrete benefit that pairing images with structured text brings to OCT classification.

Furthermore, as part of our ablation studies, we evaluated a general purpose VLM, PaLI-Gemma 2 (3B parameters), which we trained on our OCT dataset to adapt it to the OCT domain [25]. In parallel, we systematically reduced the number of training examples for our model to assess data efficiency, analyzing the impact on clinical summary quality across varying data scales (Figure 6).

### Quantitative Evaluation

We have defined two distinct tasks to showcase the models ability in providing retinal layer description and disease classification.

#### Task 1: Retinal Layer Description

In task 1, each model’s ability to generate precise, layer specific anatomical descriptions is assessed via a suite of automated language metrics reported in the upper panel of Table 1. The results demonstrate that LO-VLM produces markedly more faithful and clinically relevant textual summaries than prior VLM approaches.

**Table 1:**
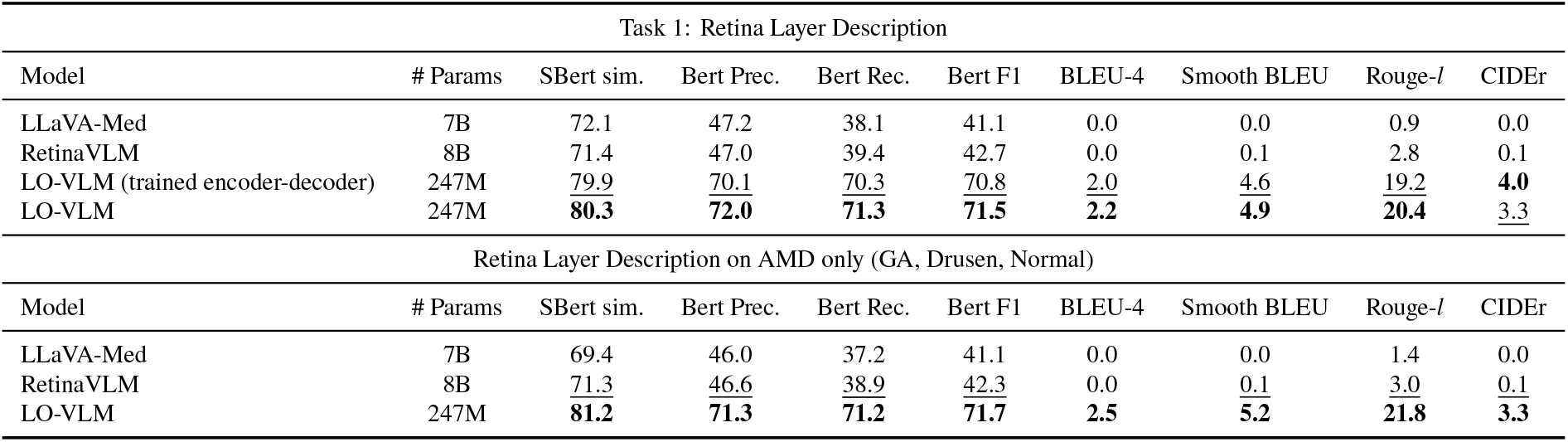
captioning performance for Task 1 (OCT layer description) across models, reported using SBERT similarity, BERTScore (Precision, Recall, F1), BLEU-4, Smoothed BLEU, ROUGE-L, and CIDEr. Highest scores are shown in bold, and second highest are underlined

| Task 1: Retina Layer Description |  |  |  |  |  |  |  |  |  |
| --- | --- | --- | --- | --- | --- | --- | --- | --- | --- |
| Model | # Params | SBert sim. | Bert Prec. | Bert Rec. | Bert F1 | BLEU-4 | Smooth BLEU | Rouge-l | CIDEr |
| LLaVA-Med | 7B | 72.1 | 47.2 | 38.1 | 41.1 | 0.0 | 0.0 | 0.9 | 0.0 |
| RetinaVLM | 8B | 71.4 | 47.0 | 39.4 | 42.7 | 0.0 | 0.1 | 2.8 | 0.1 |
| LO-VLM (trained encoder-decoder) | 247M | <u>79.9</u> | <u>70.1</u> | <u>70.3</u> | <u>70.8</u> | <u>2.0</u> | <u>4.6</u> | <u>19.2</u> | **4.0** |
| LO-VLM | 247M | **80.3** | **72.0** | **71.3** | **71.5** | **2.2** | **4.9** | **20.4** | <u>3.3</u> |
| Retina Layer Description on AMD only (GA, Drusen, Normal) |  |  |  |  |  |  |  |  |  |
| Model | # Params | SBert sim. | Bert Prec. | Bert Rec. | Bert F1 | BLEU-4 | Smooth BLEU | Rouge-l | CIDEr |
| LLaVA-Med | 7B | 69.4 | 46.0 | 37.2 | 41.1 | 0.0 | 0.0 | 1.4 | 0.0 |
| RetinaVLM | 8B | <u>71.3</u> | <u>46.6</u> | <u>38.9</u> | <u>42.3</u> | 0.0 | <u>0.1</u> | <u>3.0</u> | <u>0.1</u> |
| LO-VLM | 247M | **81.2** | **71.3** | **71.2** | **71.7** | **2.5** | **5.2** | **21.8** | **3.3** |

#### Task 2 - Disease Classification

Task 2 benchmarks two large medical VLMs and a vision only ViT-Base model on retinal disease multi-class classification against LO-VLM and its variant (trained encoder-decoder). Table 2 (lower panel) details the classification metrics for each method.

**Table 2:** Disease classification performance (Task 2) across vision-language and vision only models, reported using accuracy, F1-score, precision, and recall. The table also includes a separate evaluation on AMD specific categories (GA, Drusen, Normal). All reported differences are statistically significant (p < 0.05).

| Task 2: Disease Classification |  |  |  |  |  |  |
| --- | --- | --- | --- | --- | --- | --- |
| Model | #Params | Captioning | Accuracy | F1-score | Precision | Recall |
| LLaVA-Med | 7B | Yes | 17% | 4% | 2% | 14.3% |
| RetinaVLM | 8B | Yes | 16% | 7% | 6% | 16% |
| VIT-base | 86M | No | 83.1% | 82.9% | 85.0% | 83.1% |
| LO-VLM (trained encoder-decoder) | 247M | Yes | 84.7% | 72.8% | 73.4% | 72.6% |
| LO-VLM | 247M | Yes | **96.0%** | **96.0%** | **96.2%** | **96.0%** |

| Disease Classification on AMD only (GA, Drusen, Normal) |  |  |  |  |  |  |
| --- | --- | --- | --- | --- | --- | --- |
| Model | #Params | Captioning | Accuracy | F1-score | Precision | Recall |
| LLaVA-Med | 7B | Yes | 28.5% | 11.8% | 8.2% | 21.4% |
| RetinaVLM | 8B | Yes | 31.1% | 32% | 32% | 31% |
| LO-VLM | 247M | Yes | **94.9%** | **96.4%** | **98.1%** | **94.9%** |

#### AMD-focused Analysis

In addition, we performed an AMD-focused analysis to conduct a fair comparison between our model and RetinaVLM (trained exclusively on AMD). We constructed a balanced test subset comprising Normal, Drusen (an early stage of AMD), and GA (a late stage of AMD) cases (166 samples per class) and re-evaluated all methods.

### Task Evaluation Metrics

To comprehensively evaluate model performance across all tasks, we employed a suite of language generation and classification metrics. For Task 1, we adopted both lexical and semantic similarity metrics to evaluate the alignment between generated outputs and reference descriptions. Prior to any metric computation, we applied a targeted cleaning procedure to both the model’s outputs and the corresponding reference annotations. Specifically, we stripped out all retina layer headings, fixed labels such as “Nerve Fiber Layer:”, “Ganglion Cell Layer:”, and so forth, because these headings are structural markers rather than descriptive content. By removing them, we prevent evaluation metrics from being overly influenced by repeated, non-informative tokens that would artificially inflate scores. To capture semantic fidelity in the clinical domain, we employed SBERT [26] similarity using the abhinand/MedEmbed-large-v0.1 model from Hugging Face, a Sentence-BERT model pre-trained on large scale biomedical corpora [27]. Leveraging a medical domain SBERT model enables more accurate embedding of domain specific terminology and contextual relationships, thereby providing a more faithful assessment of semantic equivalence than general purpose models.

We further report BERT based precision, recall, and F1-score [28], which measure contextual token-level alignment using deep transformer embeddings. These metrics are particularly useful in recognizing valid paraphrasing and terminological variation that traditional n-gram based methods may overlook. To assess surface level textual similarity, we also include BLEU [29], Smooth BLEU [30], ROUGE-L [31], and CIDEr [32] scores. BLEU and its smoothed variant evaluate n-gram precision, while ROUGE-L measures the longest common subsequence, capturing both content overlap and sequential alignment. Although CIDEr was originally developed for image captioning tasks with multiple references, we apply it here in a single reference setting. In this case, CIDEr behaves similarly to BLEU but incorporates TF-IDF weighting to emphasize rare, informative n-grams. This weighting scheme provides a more nuanced assessment of content relevance, particularly beneficial in medical contexts where specific terminology carries high informational value.

For Task 2 and AMD-focused analysis, we report accuracy, precision, recall, and F1-score, which are standard metrics for multi-class classification. These collectively assess the model’s overall correctness, its ability to identify true positive cases, its sensitivity to relevant instances, and the harmonic balance between precision and recall.

### Qualitative Evaluation

We randomly selected 100 OCT images from our held-out test set, preserving the same class distribution as in our training data, and conducted a blinded, expert grading study against RetinaVLM, the only existing OCT specialized VLM. Three retina specialists independently reviewed the layer specific descriptions and diagnostic predictions produced by each model. Each grader scored every response on a 0–10 scale, with mean ± SD, median, and interquartile range reported in Table 3, according to three criteria: (i) Correctness: accuracy of the findings and diagnosis, identifying the right pathologies without errors; (ii) Completeness: inclusion of all relevant abnormalities, covering every key layer and feature; and (iii) Clarity: readability and coherence using precise, logical language that’s easy for specialists to follow.

**Table 3:**
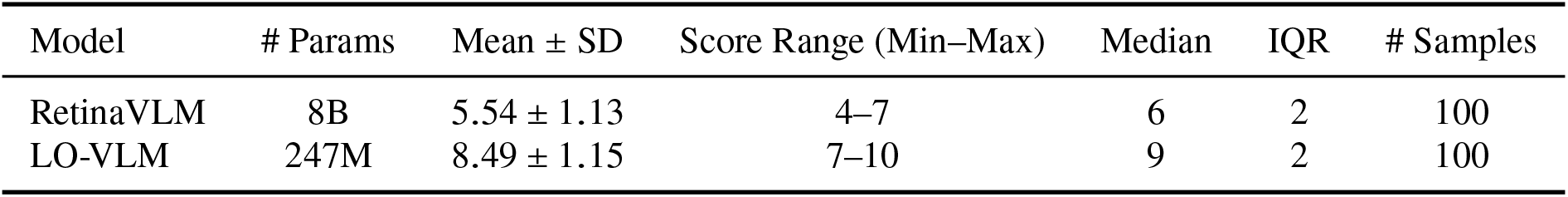
Qualitative Assessment of OCT VLMs Based on Physician Ratings (0–10 Scale). The difference between models was statistically significant (paired t-test, P < 0.0001).

### Attention-Based Interpretability

Figure 3 overlays region level attention contours for frequent anatomical keywords onto representative B-scans (more details in section). These visualizations validates model’s capacity to localize textually relevant structures and thus provide transparent, image-grounded explanations.

**Figure 3:**
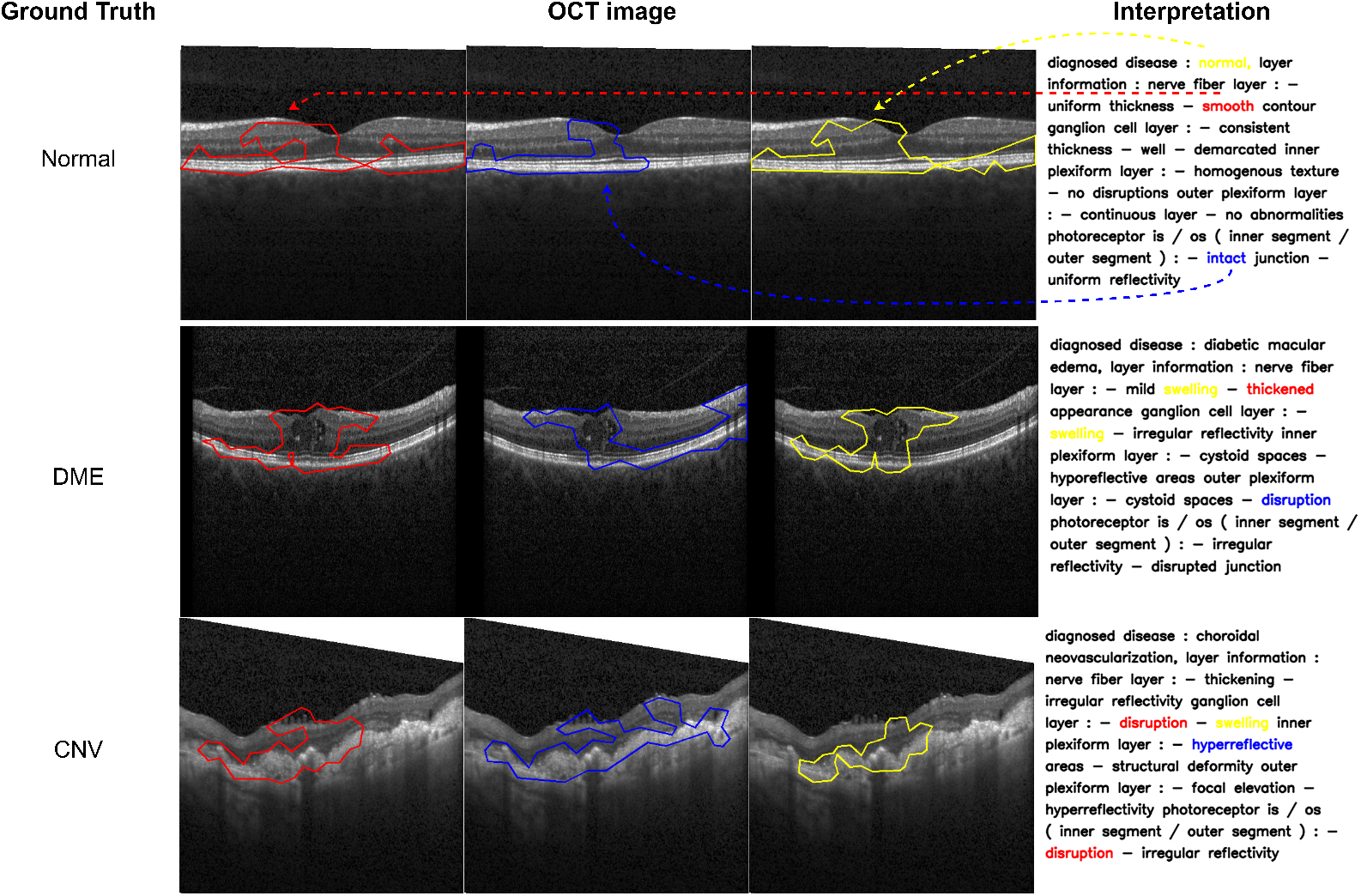
Regions contoured by LO-VLM with respect to frequent words repeated in the clinical summary dataset.

### Enhancing Readability with EYE-Llama

In clinical practice and previous ophthalmology studies, narrative paragraphs have been widely adopted because they enhance readability and conform to established reporting conventions. Although LO-VLM’s raw layer-wise outputs exhibit superior diagnostic accuracy, their tabular format may be less accessible to clinicians. To integrate the diagnostic precision of LO-VLM with the narrative clarity of traditional reporting, we apply EYE-Llama,a domain-specialized, instruction-tuned Llama 2 model, to transform structured outputs into concise, clinically accurate paragraphs [33]. Given the structured layer descriptions and diagnostic labels produced by LO-VLM, we invoke EYE-Llama with the following prompt template:

*“You are an expert ophthalmologist. Given the structured findings below, compose a concise, clinically accurate paragraph describing an OCT image suitable for a specialist audience*.

*{Retina layer-wise information}*

*please write a cohesive paragraph integrating these points without adding any new information*.*”*

To verify that this rewriting step did not introduce or omit information, we randomly audited 100 LO-VLM outputs after EYE-Llama reformatted them. A non-expert rater compared each paragraph to the original layer-wise list and found that 87 of 100 contained no added or removed clinical facts (95% CI: 0.79–0.93).

Transforming LO-VLM’s structured outputs into natural-language summaries via EYE-Llama (Figure 4) yields a cohesive narrative that improves readability, retains diagnostic precision through ophthalmology-specific terminology, and produces paragraph-formatted text ready for direct inclusion in clinical records, scholarly manuscripts, or patient-education materials.

**Figure 4:**
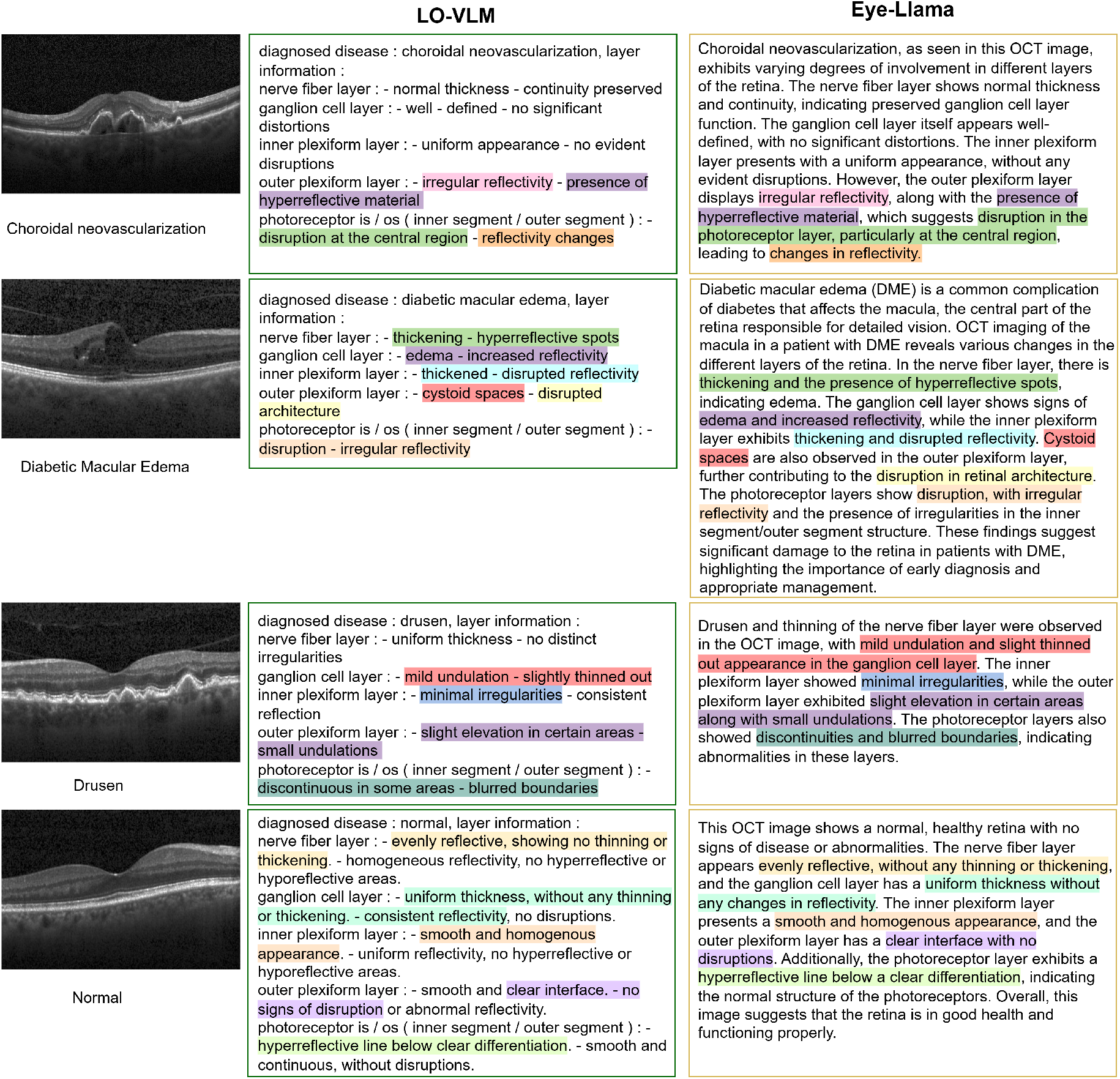
EYE-Llama enhances readability by converting LO-VLM’s structured layer-wise outputs into coherent, clinically formatted summaries.

## Ablation Studies

### Domain Specific Pre-training Investigation

We investigated whether separate pre-training of vision and text components on domain specific but unpaired data could improve performance. The ViT encoder was pre-trained on 172,000 unlabeled OCT scans using self supervised learning objectives, using masked image modeling (MIM) [34]. In MIM, a random subset of image patches is masked out during training and the network must reconstruct the missing regions from the visible context. This forces the model to learn richer, high level features of retinal structure and layer boundaries. By predicting masked regions based solely on the surrounding pixels, the ViT encoder develops an implicit understanding of OCT specific image statistics and anatomical priors. We randomly masked 60% of each OCT B-scan (preserving the top and bottom 20%) so that the model’s reconstruction loss is driven almost entirely by the central region, where the stratified retinal layers (e.g., nerve fiber, inner plexiform, and photoreceptor layers) are most clearly delineated.

In addition to the vision encoder, we pre-trained text decoder [35] on 765,000 ophthalmology text documents to capture domain specific terminology and linguistic patterns relevant to retinal disease descriptions in the language part of the VLM. This approach aimed to leverage the substantial amounts of unimodal medical data available in the public domain to enhance model understanding of OCT imaging characteristics and ophthalmic terminology. Following independent pre-training, the vision and text components were integrated and trained on our paired dataset using identical training configurations as the baseline models (Figure 5). Through evaluation of the results and comparison between LO-VLM (trained encoder-decoder) and LO-VLM (Table 1), we observe that staged unimodal adaptation fails to enrich clinical feature representations or improve performance on either descriptive or classification tasks. Instead, LO-VLM (trained encoder-decoder) requires substantially more training time and large domain specific datasets without delivering measurable gains. In contrast, direct end-to-end multimodal training of LO-VLM on paired OCT image–text examples consistently yields superior results, confirming that joint cross-modal optimization is both the most effective and the most efficient strategy for clinical OCT interpretation.

**Figure 5:**
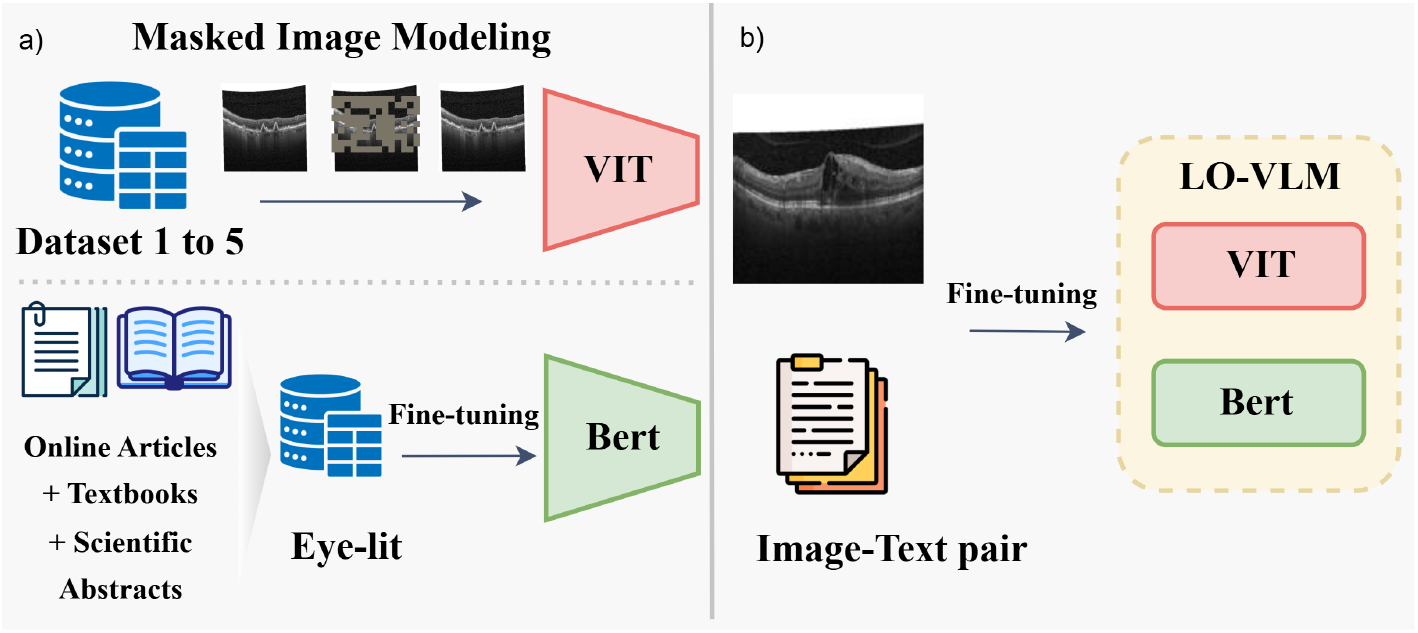
(a) Self-supervised pre-training. Top: masked image modeling is applied to five unlabeled OCT datasets to train a Vision Transformer (ViT) backbone. Bottom: the Eye-lit corpus—comprising online articles, textbooks and scientific abstracts—is used to train a BERT language model for domain specific text representations. (b) Multimodal training. The pretrained ViT and BERT are jointly tuned on paired OCT image–text examples to produce the LO-VLM model, enabling unified visual and textual understanding of retinal scans.

### Data Efficiency Analysis

We conducted systematic experiments to determine the minimum data requirements for effective OCT adaptation. Training datasets ranged from 100 to 39,000 image-text pairs, this comprehensive scaling analysis enables precise characterization of data efficiency and identifies optimal training set sizes for different deployment scenarios where data availability may be constrained.

To establish comprehensive benchmarks for data efficiency, we evaluated BLIP and PaLI-Gemma, a larger scale model trained via Low-Rank Adaptation (LoRA) [36]. For PaLI-Gemma, only 267 million of its 3 billion parameters were trained by inserting low rank decomposition matrices into the attention layers while freezing the remaining weights. This approach closely matches BLIP’s parameter count to enable a fair comparison and to assess whether the larger model can better cope with limited training data [36]. Figure 6a and 6b illustrate the performance trajectory of LO-VLM across increasing training set sizes (100-39,000 image–text pairs) over 50 training epochs (more metrics are provided in Appendix A).

**Figure 6:**
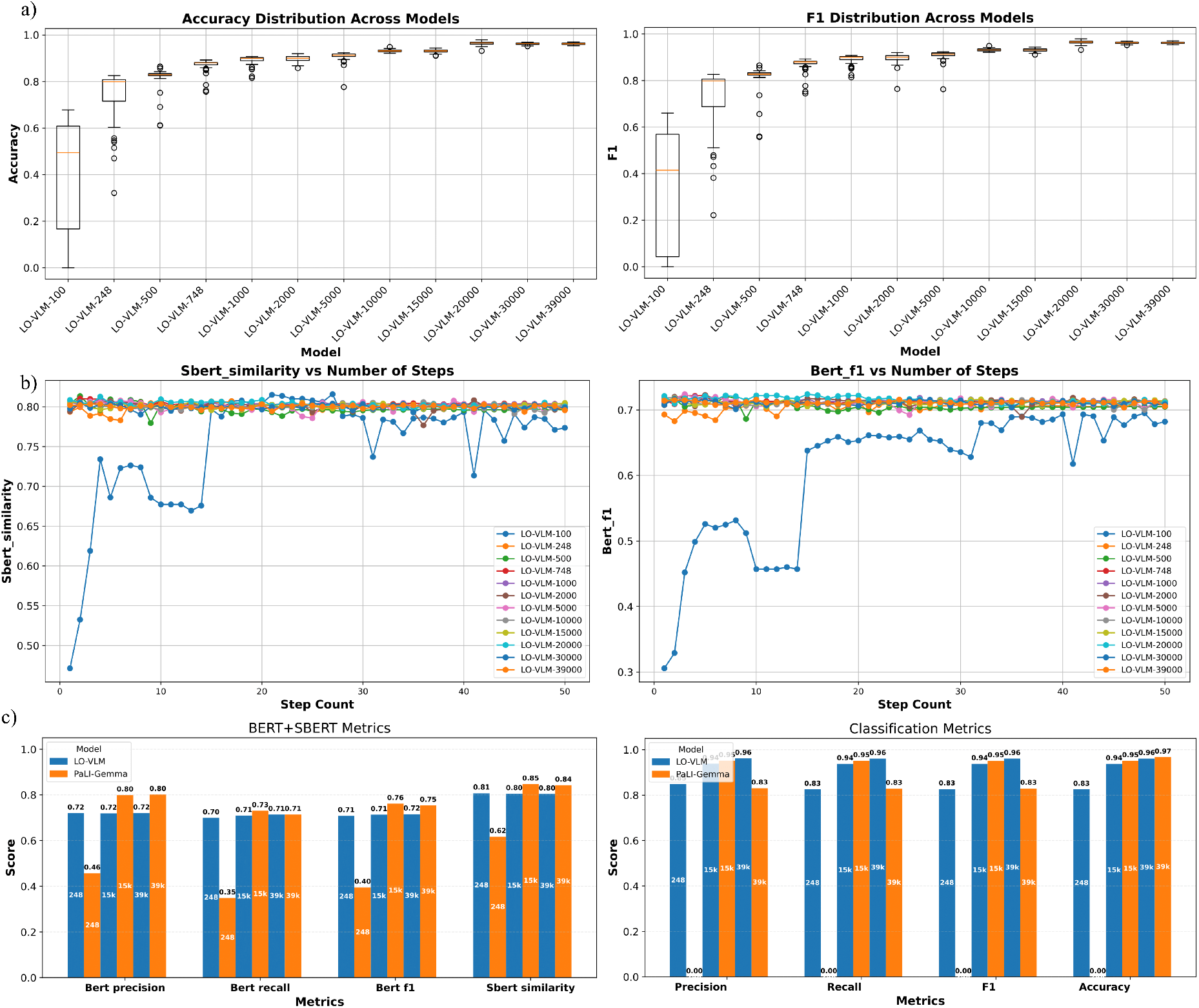
Evaluation of LO-VLM model performance. (a) Disease classification performance (F1 and Accuracy) across training steps. (b) captioning quality (BERT-F1 and SBERT similarity) across different amounts of input data. (c) Performance of LO-VLM vs. PaLI-Gemma across sample sizes (248, 15,000, 39,000).

## Discussion

Most existing VLMs are trained on general-purpose datasets such as MSCOCO and Flickr30k, which emphasize natural scenes and everyday objects and therefore lack the capacity to represent submicron retinal microstructures or to articulate pathological features in precise ophthalmic terminology. Although several medical VLMs have emerged, they are designed to span multiple imaging modalities rather than specialize in OCT. For instance, LLaVA-Med, a 7B parameter medical VLM that includes OCT among its training modalities, yields only marginal classification improvements (17% and 28%) and records an SBERT similarity of 72.1% alongside a BERTScore F1 of 41.1%. Several ophthalmology-specific VLMs have been proposed. VisionUnite [3] and Flair [37], for instance, focus exclusively on fundus imaging and therefore cannot exploit the volumetric, layer-specific information intrinsic to OCT. While RetinaVLM is currently the only model specifically designed for free form OCT summary generation, incorporating an 8 billion parameter backbone for AMD staging, referral support, and biomarker validation. However, it achieves only 16% accuracy on our six class evaluation, reflecting its limited pathology diversity. To ensure a fair comparison, we restricted the test set to the three AMD related categories (Drusen, GA, and Normal), under which RetinaVLM’s accuracy improves to 32%. Nevertheless, it still falls significantly short of LO-VLM, which maintains approximately 96% accuracy on both the full six class task and the AMD only subset. Moreover, RetinaVLM’s generated summaries exhibit lower textual alignment, with an SBERT similarity of 71.4% and a BERTScore F1 of 42.7%, underscoring its limited generalizability, a common challenge among medical models Other OCT-focused approaches include EyeClip [38] and MM-Retinal [39], which have been trained on paired OCT image–text data but do not support unconstrained clinical summary generation; notably, only GCS-M3VLT [40] has withheld its code and pretrained weights, precluding independent replication. These findings underscore three core limitations of existing medical VLMs: their poor generalizability due to limited exposure to diverse retinal conditions during training, their inability to generate free-form OCT interpretations, and their reliance on excessively large parameter counts that demand substantial computational resources. These constraints emphasize the need for efficient, OCT specific VLMs that can produce clinically meaningful outputs across a wide range of pathologies.

In this study, we conduct comprehensive assessments using both quantitative and qualitative evaluations. For Task 1 (Retinal Layer Description) in our qualitative evaluation, a blinded review was performed in which three board-certified retina specialists independently assessed anonymized OCT summaries generated by LO-VLM and RetinaVLM. LO-VLM’s outputs received a mean score of 8.5 ± 0.7 compared to 5.4 ± 0.9 for RetinaVLM (p < 0.0001), underscoring clinicians’ strong preference for LO-VLM’s anatomically grounded narratives. In our quantitative evaluation, LO-VLM outperformed LLaVA-Med and RetinaVLM across all metrics, including SBERT, BERTScore, BLEU, Smooth BLEU, ROUGE-L, and CIDEr, in both the six-class setting and the AMD-only subset (see Table 1). For Task 2 (Disease Classification), our model again outperforms all baseline VLMs in identifying retinal diseases. In this setting, we also compare our model against its own vision only component using a ViT-Base backbone with a classification head. In the six-class evaluation, LO-VLM achieved 96% accuracy (F1 = 96%), whereas the vision only ViT-Base baseline reached 83.1% accuracy (F1 = 82.9%). This performance gain is attributed to anatomical priors that guide the model’s attention toward clinically salient regions, such as drusen within the outer plexiform layer and subretinal fluid at the photoreceptor interface, thereby reducing misclassification of subtle pathological manifestations. We further compared data efficiency between LO-VLM and the PaLI-Gemma model. When trained with only 248 examples, the 3B-parameter PaLI-Gemma model failed entirely at classification, achieving 0% across all classification metrics, and underperformed in captioning with a BERTScore F1 of just 46%. In contrast, LO-VLM achieved 83% across classification metrics and a BERTScore F1 of 72% under the same low-data regime. While PaLI-Gemma’s performance improved with more data, these results highlight the superior data efficiency of LO-VLM and the importance of task-specific inductive bias. Beyond these performance metrics, the pronounced data efficiency of LO-VLM offers further insight into its practical utility. Achieving near-peak classification and summary fidelity with only 248 paired examples suggests that anatomical prompts function as potent inductive biases, effectively substituting for large volumes of raw training data. Figure 6.C illustrates how LO-VLM’s accuracy and language similarity metrics rapidly approach saturation, with diminishing gains beyond 15,000 examples. This plateau informs annotation strategy: rather than scaling purely by quantity, future efforts should prioritize diversity capturing rare pathologies, varied device settings, and demographic heterogeneity to further bolster generalizability. Moreover, the low data regime success opens the door to cost-effective deployment in resource-limited clinics and for less common disease subtypes, where large annotated corpora are infeasible. Active learning techniques could be integrated to identify the most informative OCT-text pairs, optimizing the annotation budget by focusing on edge cases and under-represented conditions.

To achieve this performance, however, we faced two primary challenges in constructing an ophthalmic VLM for OCT interpretation. The first challenge is data scarcity. Large scale paired OCT image and text corpora, where each OCT volume is linked to an expert written description, are not publicly available. Extensive repositories of OCT images and ophthalmology literature exist independently, but the paucity of curated multimodal datasets constrains the development of models that can reason over both visual and linguistic modalities. The second challenge is that existing VLMs are large scale, general-purpose models that lack the anatomical specificity needed for OCT interpretation and impose computational demands that make them impractical for clinical use. In this study, we address this gap by introducing a compact, OCT-specialized VLM with the following contributions: (i) Anatomy guided prompt integration (ii) Targeted adaptation outperforms specialized VLMs (iii) Ablation of unimodal pre-training versus end-to-end alignment (iv) Data efficiency scaling analysis (v) Clinical validation with seamless report generation.

Leveraging structured, layer specific anatomical cues, LO-VLM injects visual representations directly into the encoder and decoder attention layers to generate detailed, clinically oriented OCT summaries grounded in retinal layer visual features. This compact, domain guided adaptation significantly outperforms much larger OCT specific and general medical VLMs in producing coherent and accurate summaries, despite RetinaVLM and LLaVA-Med possessing billions of parameters. Through ablation experiments, we demonstrate that a unified, end-to-end cross-modal alignment on paired OCT image-text data consistently yields higher summary fidelity and interpretive clarity than a two stage pipeline where independent vision and language pre-training introduces additional computational overhead without commensurate benefits (Table 2, Table 1).

Despite these advances, our study has several limitations. Our dataset, although extensive, originates from a limited number of institutions and may not capture the full variability of OCT devices, scanning protocols, or rare pathologies encountered in broader clinical practice. Moreover, our evaluation was confined to six disease categories and five retinal layers, leaving finer pathological subtypes and alternative OCT modalities such as OCT angiography or longitudinal volumetric analyses unexplored. Future work should integrate real world clinical documentation and comprehensive patient metadata to enrich contextual understanding, and enlist a larger, more diverse cohort of human evaluators to rigorously assess generalizability and interpretability across varied clinical settings. Finally, extending the LO-VLM framework to incorporate complementary ophthalmic imaging modalities, including fundus photography and OCT angiography, would enable simultaneous assessment of structural and vascular biomarkers, thereby enhancing diagnostic and prognostic performance.

The development of automated OCT interpretation tools raises important ethical considerations, even in purely research contexts. This work is intended solely for research and is not designed for clinical deployment. Protecting patient privacy remains essential, requiring rigorous de-identification protocols and strong data governance. Moreover, in discussing potential applications, it is important to emphasize that such systems are intended to assist clinical expertise rather than replace it. Any future use would require careful evaluation, training, and oversight.

This work represents a step toward building clinically meaningful VLMs tailored to the unique demands of ophthalmic imaging. By focusing on anatomical grounding, interpretability, and efficiency, LO-VLM demonstrates how targeted model design can overcome the limitations of large scale general-purpose architectures in specialized medical domains. As research in medical VLMs continues to evolve, bridging the gap between domain specific insight and scalable learning frameworks will be key to advancing AI systems that are not only performant, but also clinically valuable, and data efficient.

## Methods

To enable anatomically grounded interpretation of OCT scans, we curated a paired image–text dataset from diverse public and private clinical repositories to train a VLM for retinal disease classification and generation of layer specific clinical summaries. The backbone of the VLM was based on the BLIP framework [33]. Comparative evaluations against a standalone ViT and clinical VLMs such as RetinaVLM [24]and LLaVA-Med [41], together with focused ablation studies, demonstrate that our multimodal VLM training approach on a tailored OCT dataset yields substantial improvements in diagnostic accuracy and model interpretability.

### Dataset Curation

For ablation analyses, two modality-specific datasets were curated: one comprising 172,000 spectral-domain OCT B-scans and the other, EYE-lit, containing approximately 766,000 textual excerpts. The OCT repository was assembled from multiple public and institutional sources, and encompasses both normal retinal anatomy and a broad spectrum of pathologic presentations. Details of the OCT datasets are presented in Table 4. The text-only corpus draws from two complementary sources: standard ophthalmology textbooks and peer-reviewed PubMed abstracts [33]. Textbook passages provide structured and didactic explanations of retinal layer anatomy, disease mechanisms, and clinical terminology, while research articles contribute original findings, detailed study results, and emerging insights into imaging biomarkers and therapeutic strategies.

**Table 4:** Overview of OCT datasets with source, proportion, and diagnostic categories.

| Dataset | Description | Type |
| --- | --- | --- |
| Wake Forest Dataset | A collection of high-resolution OCT images from patients with GA, classified into central, non-central GA, and no GA, including images from multiple visits. | Private |
| UIC Dataset | OCT images categorized into Control, Mild, Moderate, and Severe DR stages. Captured using an Optovue system with 3-mm, 6-mm, and 8-mm fields-of-view. | Private |
| Waterloo Dataset [42] | Over 500 high-resolution OCT images including Normal, Macular Hole, AMD, and Central Serous Retinopathy. | Public |
| Srinivasan Dataset [43] | High-resolution OCT scans from patients with DME and AMD. | Public |
| OCTDL Dataset [44] | More than 2,000 high-resolution OCT images encompassing AMD, DME, Epiretinal Membrane, Retinal Artery Occlusion, Retinal Vein Occlusion, and VMI Disease. | Public |
| Kermany Dataset [11] | 207,130 validated OCT images, categorized into CNV, DME, Drusen, and Normal. | Public |

### Image-Text Pair Dataset

The image–text pair dataset was curated by assembling 40,000 OCT images from three sources: 27,000 scans from the public Kermany et al. dataset [11], 7,200 scans from an institutional UIC collection, and 5,700 scans from a Wake Forest cohort (Figure 7.

**Figure 7:**
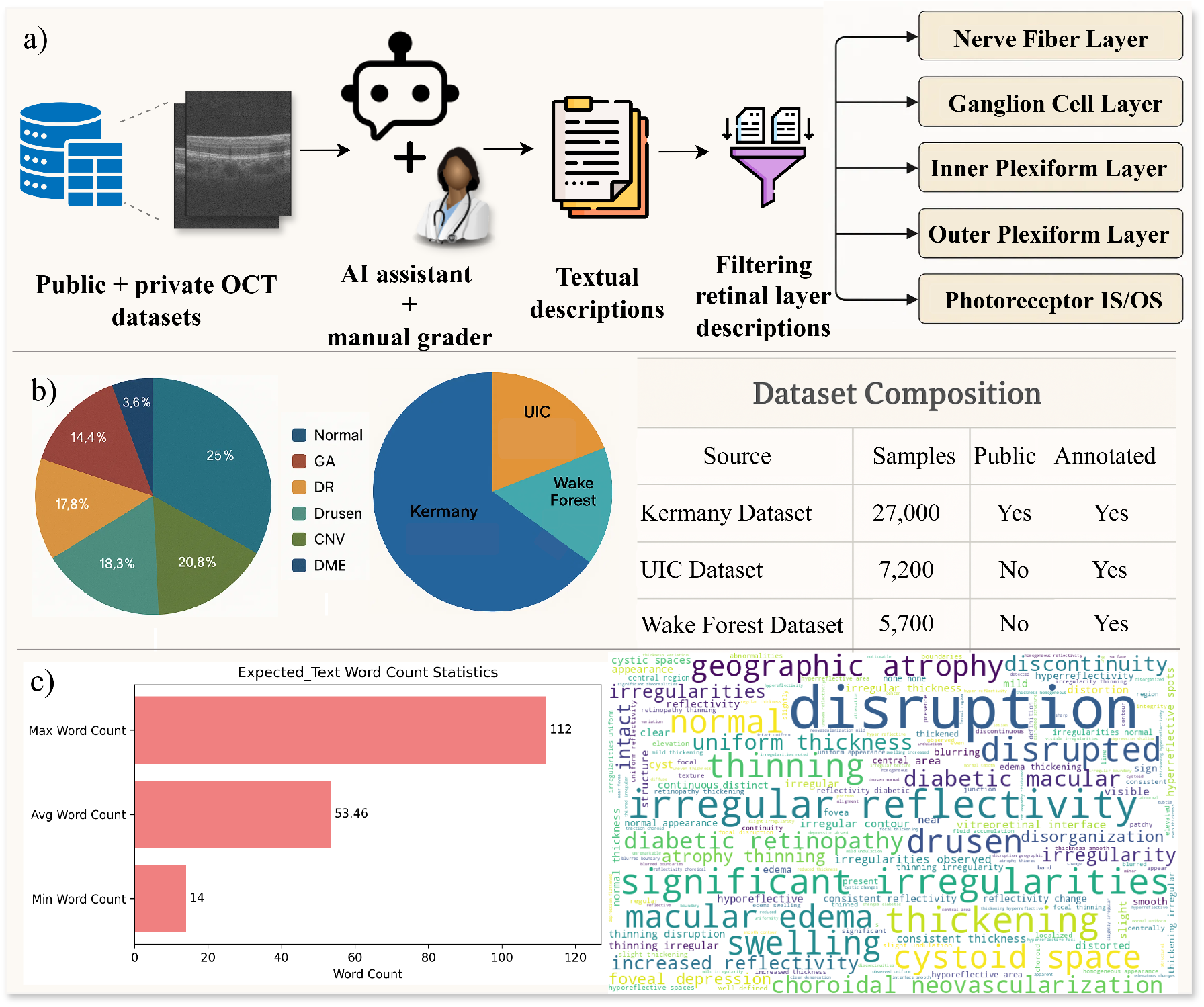
Overview of dataset pipeline and composition. In (a), raw OCT B-scans are processed by an AI assistant to generate free-text descriptions of pathologies found in OCT volume data, which are then reviewed by a clinician and assigned to five retinal layers. (b) illustrates the distribution of OCT scans by diagnostic category and by data source. The accompanying table summarizes the number of samples, public availability, and annotation status for each dataset. (c) summarizes the word count statistics of the descriptions and presents a word cloud of the most frequently used terms, emphasizing key retinal features.

Structured textual annotations were generated by prompting GPT-4o to describe each retinal layer [45](see Appendix B); these outputs were then reviewed by a retinal specialist to correct overgeneralizations and anatomical inaccuracies and to enforce consistent terminology. From the full set of OCT identifiable layers, five were selected—nerve fiber layer (NFL), ganglion cell layer (GCL), inner plexiform layer (IPL), outer plexiform layer (OPL), and photoreceptor inner/outer segment (IS/OS) and additional information of other layers if needed. These five layers consistently exhibit key diagnostic markers, such as subretinal fluid at the IS/OS interface in CNV and drusen deposits in the OPL, whereas layers beyond this set provide limited discriminative value for diagnosing the diseases in our datasets which are CNV, drusen, DR, GA, DME and healthy (normal) cases (training and testing split of 39,000 and 1,000), maintaining identical category proportions across both subsets. Each prompt concatenated the diagnostic label with these five layer specific descriptors, ensuring that annotations explicitly referenced anatomically relevant regions. This rigorously structured, multimodal curation of OCT images and layer aware text provided the foundation for multimodal training, allowing the model to learn both global diagnostic cues and localized, layer specific features essential for accurate retinal disease characterization.

### Model Architecture and Training

LO-VLM builds upon the BLIP architecture, which integrates a ViT encoder with a Transformer based text encoder–decoder framework. In this setup, the ViT processes the input image as a grid of patch tokens to generate a high dimensional visual embedding. Simultaneously, a text decoder produces language outputs conditioned on these visual features. During training, BLIP is optimized using three complementary objectives:

- Image–Text Matching (ITM): A cross-modal Transformer block incorporating bi-directional self-attention, cross-attention, and feed-forward layers is trained to classify whether a given image–text pair is semantically aligned.
- Image–Text Contrastive (ITC): A dual encoder contrastive loss function draws matched image and text embeddings closer within a shared latent space while repelling mismatched pairs.
- Image-Grounded Language Modeling (LM): A causal language modeling objective trains the decoder to auto-regressive generate captions, token-by-token, based on the encoded visual representation.

This modular architecture supports both retrieval based tasks (via ITM and ITC) and generative captioning (via LM), as illustrated in figure 1.

We trained our VLM to translate OCT B-scans into anatomically precise, layer-aware retinal descriptions by using structured prompts that combined each scan’s diagnostic label (e.g., “CNV,” “Drusen”) with its five layer-specific descriptors (NFL, GCL, IPL, OPL, IS/OS). Each training instance comprises a grayscale OCT scan, its segmentation map, and a text prompt formatted as: “<LayerName>: clinical summary of observed pathology or normal anatomy.” This formulation injects anatomical priors into the learning process (Fig. 1). In each update, a given segmented B-scan is first processed by a ViT encoder to obtain a dense visual embedding encapsulating both global retinal morphology and subtle layer-wise reflectivity patterns. This embedding is shared across the original training objectives to generate detailed, anatomically grounded summaries such as “IS/OS: intact, continuous” or “IPL: focal hyperreflectivity indicating edema.”

The Retina Layer–Text Matching (RLTM) objective assesses whether each scan–summary pair is correctly aligned, with positive examples linking scans to their respective layer descriptions. This promotes fine-grained visual–textual alignment at the anatomical layer level. The Retina-Layer Contrastive Alignment (RLCA) objective further refines this alignment by drawing matched scan–summary embeddings closer in a shared latent space while separating mismatched pairs, thereby guiding the encoder toward anatomically salient features.

Finally, the Layer-Conditioned Language Modeling (LCLM) objective conditions captioning on the visual representation of each segmented scan, encouraging the decoder to produce clinically precise, anatomically grounded summaries initiated by the corresponding layer identifier.

For training the LO-VLM model, we followed the hyperparameter configuration outlined in the original baseline framework, with targeted adjustments. A batch size of 32 was selected as the maximum capacity supported by our 49GB GPU, and the model was trained for 50 epochs. For captionign task, the maximum output length was set to 256 tokens, which was empirically sufficient to capture detailed, layer-wise retinal descriptions. The training dataset consisted of 39,000 OCT scan–summary pairs, with 1,000 samples reserved for testing. This configuration enabled the development of a clinically grounded framework capable of producing anatomically precise summaries from OCT B-scans.

### Vision Only Baseline: ViT-Base Training on OCT Images

VLMs have recently gained attention for their ability to generate descriptive reports and support decision making based on visual data. To benchmark this new paradigm against traditional image classification, we trained a standalone ViT-Base model solely on OCT scans, 39,000 images for training and 1,000 for evaluation, excluding any accompanying text information. Each B-scan was uniformly resized to 224 × 224 pixels, and the pretrained 768-dimensional classifier head was replaced with a linear layer projecting to our six disease categories. The network was optimized with cross-entropy loss using the Adam algorithm (learning rate =1 × 10^−5^), a batch size of 32, and a 50 epoch schedule. This unimodal baseline establishes a clear benchmark for quantifying the added value of language grounding and cross-modal alignment in our LO-VLM system. ViT-Base was selected to mirror the vision encoder of our LO-VLM framework, ensuring architectural consistency so that any performance differences can be attributed solely to the addition of textual information. As a widely adopted transformer based backbone, ViT-Base benefits from extensive pretraining on large scale image corpora and has proven effective when trained on medical imaging tasks, including OCT classification [9, 46].

### Specialized and Foundation VLMs

We compared our results to RetinaVLM which is the only VLM specifically trained for OCT interpretation. This baseline employs an 8B parameter Llama 3 backbone paired with a ResNet vision encoder and was originally trained on a proprietary OCT dataset that was subsequently extended using an LLM assistant; this augmented dataset was then used to train the model. Beyond this specialized baseline, we evaluated a medical Model, LLaVA-Med (7B params). LLaVA-Med is a medical image–captioning model trained on diverse imaging modalities, including OCT. The trained “microsoft/llava-med-v1.5-mistral-7b” model was retrieved from the Hugging Face repository and its performance was assessed using the developer recommended Mistral Instruct conversation template. In accordance with the original RetinaVLM publication, we then employed the patient-triage prompt that most closely aligns with RetinaVLM’s layer-wise summary (Figure 2 more samples are provided in Appendix C).

### Token Specific Gradient Saliency for Anatomical Mapping

We integrate gradient-based saliency to elucidate how each descriptive term in our OCT clinical summary maps to distinct retinal microstructures. By deriving token-specific saliency maps, we confirm that semantic labels correspond to anatomically relevant regions, thereby strengthening interpretability and fostering clinical trust. Gradient-based saliency provides a rigorous mechanism for attributing individual clinical summary tokens to precise image regions. For each keyword generated by the VLM, we backpropagate its output logit through the vision encoder (w.r.t. first LayerNorm of the last transformer block) and capture the resulting gradients at an intermediate normalization layer (Figure 3). Summing the absolute values of these gradients across feature channels yields a coarse saliency grid, which is interpolated to the original image resolution and thresholded to isolate the most influential pixels. This token-wise mapping not only offers deep insight into the model’s decision-making process but also verifies that its semantic outputs are firmly grounded in clinically meaningful features[47, 48].

### Statistical analysis

Automatic per-sample metrics (e.g., per-image correctness for classification, SBERT similarity, BERTScore for text quality) were computed for each test image (*N* = 10) for every model. Each metric was then averaged across the test set to obtain a single summary value per model (reported as means). All models were evaluated on the same held-out images to ensure a like-for-like comparison.

#### Human evaluation

For human evaluation (*number of samples* = 100 OCT images; 3 graders), each grader scored every generated summary using the same rubric. Scores were averaged across graders for each sample, and these per-sample averages were summarized as mean ± SD across the 100 clinical summaries.

#### Hallucination audit

We sampled 100 LO-VLM→EYE-Llama narratives and manually checked whether any facts were added or removed during paragraph reformatting. In 87 of 100 cases, no factual changes were observed (exact binomial 95% CI: 0.79–0.93).

#### Software

Analyses were performed in Python (numpy/pandas, scipy, matplotlib, seaborn libraries).

## Supporting information

Supplemental Doc

## Data availability

The curated layer-specific OCT image–text dataset comprises 40 000 paired B-scans and descriptions assembled from three sources: 27 000 scans from the public Kermany dataset, 7 200 scans from an institutional UIC collection, and 5 700 scans from a Wake Forest cohort. Public OCT repositories—Kermany et al. [11], OCTID [42], Srinivasan et al. [43], and OCTDL [44]—are available at their original archives. Private UIC and Wake Forest datasets cannot be open sourced. The public part of the image-text dataset consisting 27k pairs, will be open sourced upon publication.

## Code availability

All custom code for dataset curation, model training, evaluation scripts, and the pretrained LO-VLM weights will be made publicly available upon publication under the MIT license.

## Acknowledgements

This work was supported by NEI R15EY035804 (MNA) and UNC Charlotte Faculty Research Grant (MNA).

## Author contributions

T.H. and M.N.A. conceived and designed the study; H.T. and M.N.A. supervised the study. T.H. and S.G. curated and annotated the dataset; J.T.S. and T.L. evaluated the results of the model; J.I.L. and A.C.T. provided clinical expertise and reviewed annotations; T.H. trained and developed the LO-VLM model; T.H. and S.G. analyzed results; T.H. drafted the manuscript; all authors reviewed and edited the manuscript.

