## Supplemental Doc for "Compact Vision–Language Models Enable Efficient and Interpretable Automated OCT Analysis Through Layer Specific Multimodal Learning"

### Supplemental Materials

#### Appendix A LO-VLM performance metrics

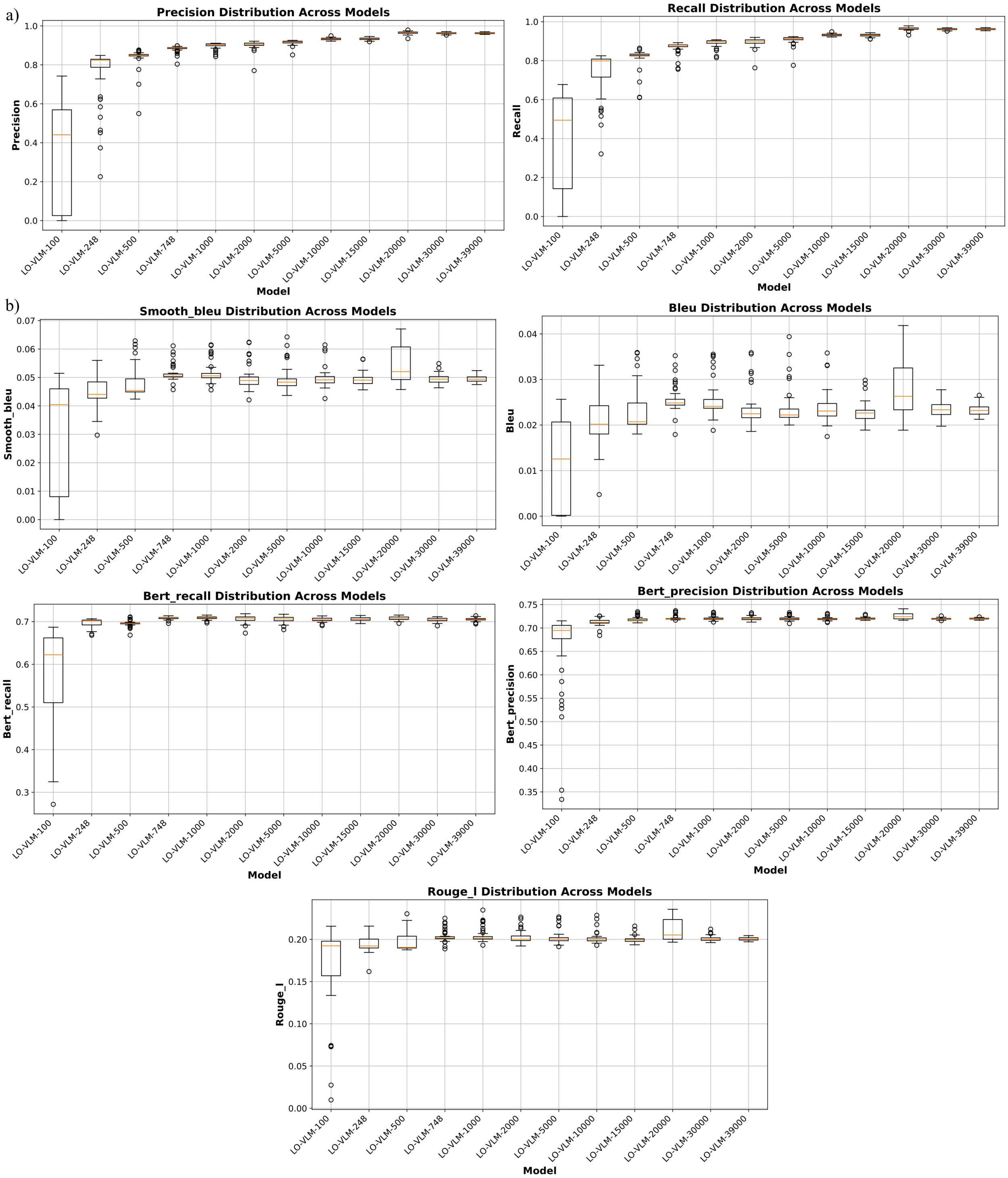


Figure S1: LO-VLM's performance across training set sizes ranging from 248 to 39,000 image–text pairs. (a) reports the classification performance metrics, while (b) presents the complementary text-generation (NLP) metrics that augment those shown in Figure 4.

### Appendix B GPT4o Prompt for Generating Layer-wise Summary

We employed the following instruction to generate clinical summaries from GPT-4o, replacing the placeholder “**” with the specific pathology depicted in each OCT image.

*"Focus on the OCT image provided and the layers you see in detail. This OCT image has been diagnosed as ** by an ophthalmologist. For each layer below, write some keywords about its irregularities in the image (be specific about what the irregularity is and where it can be seen in the image).*

*We also want to consider the vitreoretinal interface above, the choroid below and foveal depression.*

*Layers:*

*### Inner Retina:*

*#### Nerve Fiber Layer:*

-

-

*#### Ganglion Cell Layer:*

-

-

*#### Inner Plexiform Layer:*

-

-

*#### Inner Nuclear Layer:*

-

-

*### Outer Retina:*

*#### Outer Plexiform Layer:*

-

-

*#### Outer Nuclear Layer:*

-

-

*#### External Limiting Membrane:*

-

-

*#### Photoreceptor IS/OS (Inner Segment/Outer Segment):*

-

-

*#### Retinal Pigment Epithelium:*

-

-

*### Additional Structures:*

*#### foveal depression:*

-

-

*#### vitreoretinal interface:*

-

-

*#### choroid:*

-

-

*After writing the above keywords for each layer. Write a comma separated list of irregularities (just write keywords) indicative of **.*

*Do not write bullet points.*

*### keywords:"*

### Appendix C Sample Generated Clinical Summaries

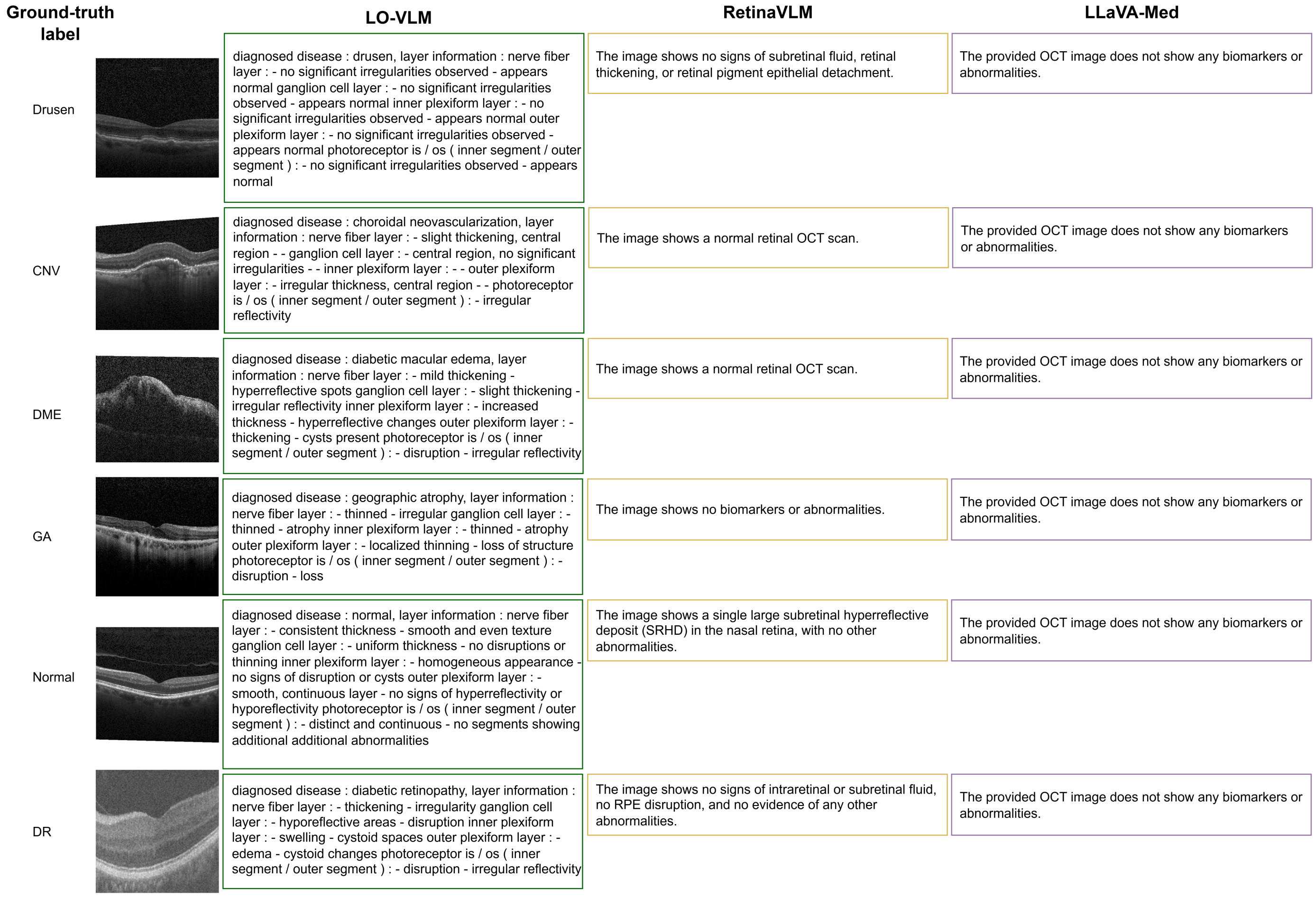


Figure S2: Qualitative comparison of OCT image interpretation across models. From left to right: (1) ground-truth disease label and exemplar B-scan, (2) LO-VLM generated layer-wise descriptions, (3) RetinaVLM outputs, and (4) LLaVA-Med outputs. Rows correspond (top to bottom) to Drusen, CNV, DME, GA, normal retina, and DR.
